# Processing of Auditory Novelty Across the Cortical Hierarchy: An Intracranial Electrophysiology Study

**DOI:** 10.1101/290106

**Authors:** Kirill V. Nourski, Mitchell Steinschneider, Ariane E. Rhone, Hiroto Kawasaki, Matthew A. Howard, Matthew I. Banks

**Author notes:** **Corresponding author:** Kirill V. Nourski, MD, PhD, Department of Neurosurgery, The University of Iowa, 200 Hawkins Dr. 1815 JCP, Iowa City, IA 52242 USA.

## Abstract

Under the predictive coding hypothesis, specific spatiotemporal patterns of cortical activation are postulated to occur during sensory processing as expectations generate feedback predictions and prediction errors generate feedforward signals. Establishing experimental evidence for this information flow within cortical hierarchy has been difficult, especially in humans, due to spatial and temporal limitations of non-invasive measures of cortical activity. This study investigated cortical responses to auditory novelty using the local/global deviant paradigm, which engages the hierarchical network underlying auditory predictive coding over short (‘local deviance’; LD) and long (‘global deviance’; GD) time scales. Electrocorticographic responses to auditory stimuli were obtained in neurosurgical patients from regions of interest (ROIs) including auditory, auditory-related and prefrontal cortex. LD and GD effects were assayed in averaged evoked potential (AEP) and high gamma (70-150 Hz) signals, the former likely dominated by local synaptic currents and the latter largely reflecting local spiking activity. AEP LD effects were distributed across all ROIs, with greatest percentage of significant sites in core and non-core auditory cortex. High gamma LD effects were localized primarily to auditory cortex in the superior temporal plane and on the lateral surface of the superior temporal gyrus (STG). LD effects exhibited progressively longer latencies in core, non-core, auditory-related and prefrontal cortices, consistent with feedforward signaling. The spatial distribution of AEP GD effects overlapped that of LD effects, but high gamma GD effects were more restricted to non-core areas. High gamma GD effects had shortest latencies in STG and preceded AEP GD effects in most ROIs. This latency profile, along with the paucity of high gamma GD effects in the superior temporal plane, suggest that the STG plays a prominent role in initiating novelty detection signals over long time scales. Thus, the data demonstrate distinct patterns of information flow in human cortex associated with auditory novelty detection over multiple time scales.

## Introduction

Far from being passive receivers of sensory information, humans are actively engaged in the process of sensation. Sensory perception and motor responses to identical stimuli can vary based on attention and expectation (den Ouden et al., 2012), but how such contextual modulation is implemented at a systems level is unclear. Under the predictive coding hypothesis (Bar, 2009; Clark, 2013; Friston, 2005), sensory predictions are generated at high levels in the cortical hierarchy and projected via feedback (FB) connections to lower levels, where they are compared with incoming sensory data. Error signals arising from violations of these expectations are then projected back to higher levels via cortical feedforward (FF) connections (Bastos et al., 2012; Mumford, 1992; Rao and Ballard, 1999). Stability of this information exchange is central to leading theories of brain function and consciousness (Dehaene and Changeux, 2011; Friston, 2010; Mashour, 2013; Tononi et al., 2016). Although there is extensive circumstantial support for the predictive coding hypothesis (Bastos et al., 2012; Heilbron and Chait, 2017), direct experimental evidence is only beginning to emerge (Blank and Davis, 2016; Egner et al., 2010). Detailed understanding of the spatial distribution, timing, and directionality of information flow during predictive sensory processing is lacking. For example, activation of higher-order cortical areas should precede that of lower-order areas in generating expectations, while prediction error signals should unfold in the opposite order. However, there has been limited investigation of the temporal sequence of activation of cortical regions during predictive coding. Even the identity of these interacting regions remains ambiguous (Deouell, 2007), and depends on the time scale over which predictive coding operates and the level of engagement of attentional networks in the task at hand (Wacongne et al., 2011).

Auditory novelty detection is a sensory task postulated to activate predictive coding networks (Garrido et al., 2009). The present study took advantage of the local/global deviant (LGD) stimulus paradigm (Bekinschtein et al., 2009). This paradigm is an elaboration of the classic oddball paradigm designed to investigate auditory novelty detection over multiple time scales and engage both pre-attentive and conscious perceptual processes (Bekinschtein et al., 2009; Strauss et al., 2015). Stimuli consist of repeating tokens, with repetition within the stimulus (“local standards”; LS) establishing expectation for the last token over short time scales (<1 s). Repeated stimulus sequences (“global standards”; GS) establish expectation on a longer time scale (5–10 s). Violations of expectation by “local deviants” (LD) and “global deviants” (GD) trigger enhanced responses that are interpreted as prediction error signals. LD effects have been associated with the scalp-recorded mismatch negativity (MMN) response (Naatanen and Alho, 1995), whose generators are reported to be localized primarily to the auditory cortex (Bekinschtein et al., 2009; Joos et al., 2014). GD effects have been associated with the P3b component of the event-related potential (ERP) (Kok, 2001), generated within a broader network that includes temporoparietal and prefrontal regions (Bekinschtein et al., 2009; Halgren et al., 1998; Smith et al., 1990).

Key questions remain about the generators of LD and GD effects. For example, although some studies localized the MMN and LD effects to auditory cortex (Alho, 1995; Bekinschtein et al., 2009; El Karoui et al., 2015), others describe a broader network including auditory cortex, surrounding temporoparietal auditory-related areas and prefrontal cortex (PFC) (Durschmid et al., 2016; Rinne et al., 2000; Schonwiesner et al., 2007). Furthermore, even basic information about the sequence of activation of these regions is unclear, with the timing of frontal relative to temporal generators of the MMN ranging from lag to lead (Rinne et al., 2000; Schonwiesner et al., 2007; Yago et al., 2001). It is well-established that GD effects manifest with longer latencies (Bekinschtein et al., 2009), suggesting greater involvement of higher order processing compared to the LD effect. Whether the response to the GD stimulus is generated first in PFC, auditory-related or auditory cortex is unclear, as is the associated timing of signal propagation up and down the cortical hierarchy. Here, we provide a detailed picture of the spatial and temporal activation patterns during auditory novelty detection over multiple time scales, taking advantage of the superior spatial and temporal resolution offered by intracranial recordings. This work builds upon previous ECoG studies of auditory novelty detection (Durschmid et al., 2016; Edwards et al., 2005; El Karoui et al., 2015) by analyzing high density ECoG data obtained simultaneously from all cortical levels envisioned to be involved in auditory predictive coding, from core (primary) auditory cortex to PFC.

## Methods

### Subjects

Study subjects were six neurosurgical patients with medically refractory epilepsy who had been implanted with intracranial ECoG electrodes to identify resectable seizure foci. Research protocols were approved by the University of Iowa Institutional Review Board and the National Institutes of Health, and written informed consent was obtained from all subjects. Research participation did not interfere with acquisition of clinically necessary data, and subjects could rescind consent for research without interrupting their clinical management. Experiments were performed when the patients returned to the operating room to undergo electrode removal and seizure focus resection surgery. The demographic and seizure focus data for each subject are presented in Table 1. Recording sites that were confirmed to be involved in seizure onset were excluded from analysis. All subjects were native English speakers, right-handed and had left language dominance as determined by Wada tests. All subjects had pure-tone thresholds within 30 dB hearing level between 125 Hz and 8 kHz (Supplementary Fig. 1). Word recognition scores, as evaluated by spondees presented via monitored live voice, were 88% or higher, and speech reception thresholds were within 20 dB in all tested subjects. Cognitive function, as determined by standard neuropsychological assessments, was in the average range in all subjects.

**Table 1.** Subject demographics.

| Subject<sup>1</sup> | Age | Sex<sup>2</sup> | Seizure focus | Surgical procedure |
| --- | --- | --- | --- | --- |
| R369 | 30 | M | R medial temporal lobe | R anterior and medial temporal lobectomy |
| L372 | 34 | M | L temporal pole | L anterior and medial temporal lobectomy |
| R376 | 48 | F | R medial temporal lobe | R anterior and medial temporal lobectomy |
| R394 | 24 | M | R amygdala | R anterior and medial temporal lobectomy |
| R399 | 22 | F | R temporal lobe with early propagation to R inferior lateral frontal lobe | R temporal lobectomy and R inferior lateral frontal lobectomy |
| L400 | 59 | F | L amygdala | L anterior and medial temporal lobectomy |

<sup>1</sup>Letter prefix of the subject code denotes the side of electrode implantation over auditory cortex, and the side of seizure focus (L = left; R = right). Most subjects had, to varying degrees, bilateral coverage of other regions of the brain.
<sup>2</sup>F = female; M = male

Subject R394 had previously undergone resection of a cavernoma in the anterior medial temporal lobe, in which part of the amygdala and head of hippocampus were removed. Adhesion of the dura mater to the cortical surface secondary to that surgery precluded subdural placement of surface electrode arrays in this subject. Instead, surface arrays were placed epidurally. While the epidural recordings had diagnostic utility for seizure focus detection, their signal-to-noise ratio and localization of electrode contacts in relation to the cortical surface anatomy were unfavorable for research data acquisition. Consequently, recordings obtained from the epidural contacts were excluded from analyses in the present study. In this subject, cortex that corresponded to all studied brain regions (see below) was spared, with the exception of planum polare. Additionally, this subject had normal hearing, normal intelligence, and exhibited above-average performance on the behavioral task. This provided a justification for inclusion of data obtained from this subject in the study.

### Stimulus and procedure

Stimulus generation was controlled by a TDT RZ2 real-time processor (Tucker-Davis Technologies, Alachua, FL). Experimental stimuli were vowels /a/ and /i/, presented in an LGD paradigm (Bekinschtein et al., 2009; El Karoui et al., 2015; Strauss et al., 2015) (Fig.1). The vowels were excised from the steady-state vowel portions of consonant-vowel stimuli /had/ and /hid/, spoken by a female (F_0_ = 232 Hz and 233 Hz, respectively) (Hillenbrand et al., 1995). On each trial, five 100 ms vowels, normalized to the same root-mean-square amplitude, gated with 5 ms on/off ramps and separated by 50 ms silent intervals, were presented, with the fifth vowel being either the same as the first four (LS) or different (LD; Fig. 1a). This difference constituted local deviance: /aaaaa/ and /iiiii/ were LS stimuli, while /aaaai/ and /iiiia/ were LD stimuli.

**Figure 1.**
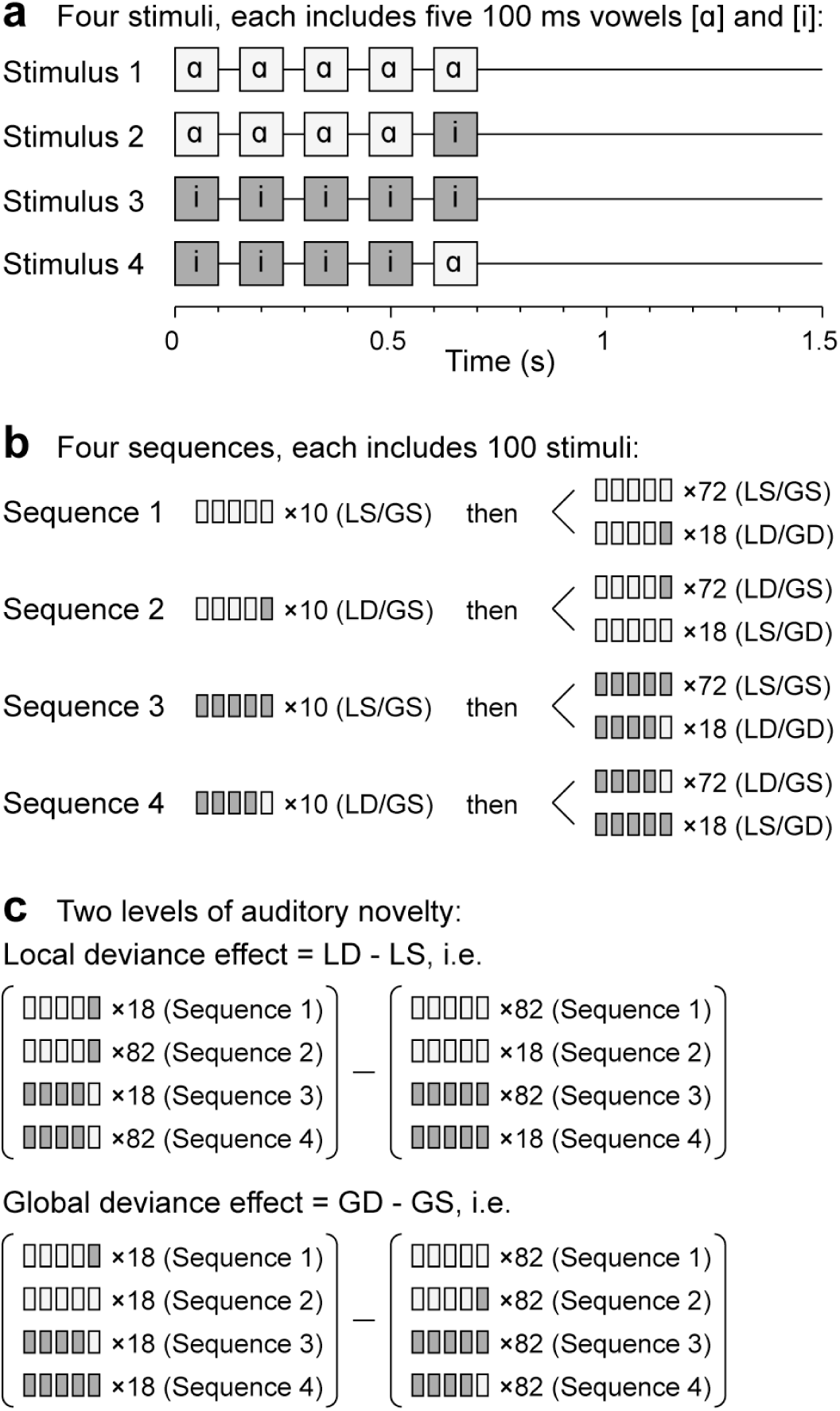
Local global deviant experimental paradigm. **a:** Schematic of the four experimental stimuli. **b:** Stimulus sequences. **c:** Comparisons between trials to characterize local and global deviance responses. Adapted from Strauss et al. (2015).

The stimuli were presented in four sequences, with the order of the sequences randomized across subjects (Fig. 1b). Each sequence began with a recorded instruction that defined the task and the target (GD) stimulus to the subject, e.g., for Sequence 2: “This time, press the button when you hear this sound: /aaaaa/. Once again, press the button when you hear this sound: /aaaaa/.” This was followed by a habituation sequence of 10 trials that established the GS stimulus (e.g. /aaaai/ for Sequence 2), and then by 72 GS trials and 18 GD trials, presented in a pseudorandom order. The difference in presentation frequency thus constituted the global deviance, and the identity of the GD stimulus changed across the four sequences within each block. This design allowed for the simultaneous evaluation of responses to auditory novelty on two different time scales (local and global; Fig. 1c). The intertrial interval varied within a Gaussian distribution (onset-to-onset mean 1500 ms, S.D. = 10 ms) to reduce heterodyning in the recordings secondary to power line noise. The duration of the experimental block was 11 minutes (400 ∼1.5 s trials and four 15 s instruction segments).

Stimuli were delivered to both ears via insert earphones (ER4B, Etymotic Research, Elk Grove Village, IL) that were integrated into custom-fit earmolds. Acoustic stimulation was performed at a comfortable level, typically 60-65 dB SPL. The target detection task was used to control for and provide measures of the subjects’ level of attention. The hand ipsilateral to the hemisphere from which recordings were made was used to operate the response button. This was done to minimize contributions of preparatory (motor planning), motor and somatosensory responses associated with the button press to recorded neural activity as opposed to auditory deviance processing *per se*.

The data presented here were collected within the context of a larger study on effects of general anesthesia on auditory cortical responses during a pre-drug baseline period (Nourski et al., 2018). As part of that study, the subjects’ overall level of alertness was evaluated over the course of the experiment using the Observer’s Assessment of Alertness/Sedation (OAA/S) scale, which ranges from OAA/S = 5 for fully awake to OAA/S = 1 for unresponsive even to noxious stimuli (Chernik et al., 1990). Because we observed that task performance even prior to administration of anesthesia was related to level of alertness, we present the task performance data along with OAA/S scores measured immediately before and after the recordings presented here.

### Recording

ECoG recordings were made using subdural and depth electrode arrays (Ad-Tech Medical, Racine, WI) placed on the basis of clinical requirements to identify seizure foci (Nagahama et al., 2017; Reddy et al., 2010). Electrode implantation, recording and ECoG data analysis have been previously described in detail (Howard et al., 1996, 2000; Nourski and Howard, 2015; Reddy et al., 2010). In brief, the subdural arrays consisted of platinum-iridium disc electrodes (2.3 mm exposed diameter, 5-10 mm inter-electrode distance) embedded in a silicon membrane. Subdural strip and grid arrays were implanted over lateral and ventral surfaces of temporal, and frontal lobe, and lateral parietal cortex. Depth electrode arrays (8-12 macro contacts, spaced 5 mm apart) targeting superior temporal plane, including Heschl’s gyrus (HG), were stereotactically implanted along the anterolateral-to-posteromedial axis of the gyrus. Additional arrays targeted insular cortex and provided coverage of posteromedial HG (HGPM), planum temporale (PT) and planum polare (PP). This configuration was used to provide a more accurate assessment of suspected temporal lobe seizure foci than could be provided subdural electrodes alone by bracketing epileptogenic zones from dorsal, ventral, medial and lateral aspects. Depth electrodes that targeted mesial temporal lobe structures (amygdala and hippocampus) provided additional coverage of auditory-related cortex within superior temporal sulcus. A subgaleal electrode was used as a reference in all subjects.

Reconstruction of the anatomical locations of implanted electrodes and their mapping onto a standardized set of coordinates across subjects was performed using FreeSurfer image analysis suite (Version 5.3; Martinos Center for Biomedical Imaging, Harvard, MA) and in-house software (see Nourski et al., 2014, for details). In brief, subjects underwent whole-brain high-resolution T1-weighted structural magnetic resonance imaging (MRI) scans (resolution and slice thickness ≤1.0 mm) before electrode implantation. After electrode implantation, subjects underwent MRI and thin-slice volumetric computerized tomography (CT) (resolution and slice thickness ≤1.0 mm) scans. Contact locations of the depth and subdural electrodes were first extracted from post-implantation MRI and CT scans, respectively. These were then projected onto preoperative MRI scans using non-linear three-dimensional thin-plate spline morphing, aided by intraoperative photographs. Data from multiple subjects were pooled by transforming the electrode locations into standard Montreal Neurological Institute (MNI) coordinates. This was done for each contact using linear coregistration to the MNI152 T1 average brain, as implemented in FMRIB Software library (Version 5.0; FMRIB Analysis Group, Oxford, UK). Left hemisphere MNI x-axis coordinates (x_MNI_) were then multiplied by (−1) to map them onto the right-hemisphere common space. Contacts were then projected onto the right lateral hemispheric surface and right superior temporal plane of the FreeSurfer average template brain.

Regions of interest (ROIs) included, in roughly ascending hierarchical order, core auditory cortex in HGPM, non-core auditory areas (anterolateral portion of Heschl’s gyrus [HGAL], PT, PP, STG), auditory-related (insular cortex, superior temporal sulcus, middle temporal gyrus [MTG], supramarginal and angular gyrus), and PFC, including inferior (IFG), middle and superior frontal gyrus, orbital gyrus and transverse frontopolar gyrus (Table 2). Assignment of recording sites to ROIs was based upon anatomical reconstructions of electrode locations in each subject. For subdural arrays, it was informed by automated parcellation of cortical gyri (Destrieux et al., 2010, 2017) as implemented in the FreeSurfer software package. For depth electrodes, ROI assignment was informed by MRI sections along sagittal, coronal and axial planes. Recording sites identified as seizure foci or characterized by excessive noise, depth electrode contacts in white matter or outside brain, and epidurally implanted arrays in subject R394 were excluded from analyses and thus are not listed in Table 2. These criteria led to exclusion of 34/249 contacts in subject R369, 33/214 in L372, 27/226 in R376, 49/74 in R394, 50/238 in R399, and 52/210 in L400.

**Table 2.**
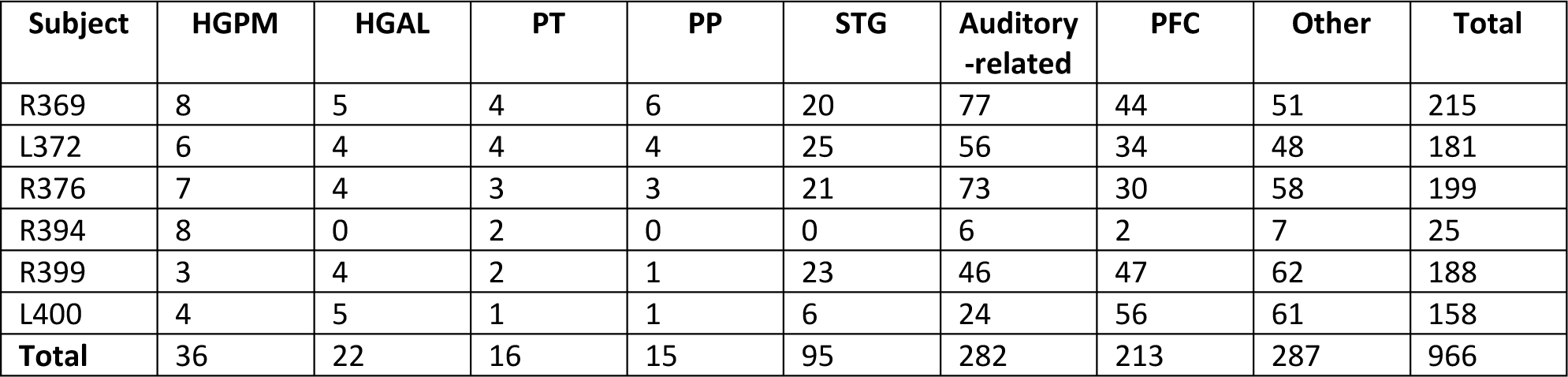
Electrode coverage.

| Subject | HGPM | HGAL | PT | PP | STG | Auditory -related | PFC | Other | Total |
| --- | --- | --- | --- | --- | --- | --- | --- | --- | --- |
| R369 | 8 | 5 | 4 | 6 | 20 | 77 | 44 | 51 | 215 |
| L372 | 6 | 4 | 4 | 4 | 25 | 56 | 34 | 48 | 181 |
| R376 | 7 | 4 | 3 | 3 | 21 | 73 | 30 | 58 | 199 |
| R394 | 8 | 0 | 2 | 0 | 0 | 6 | 2 | 7 | 25 |
| R399 | 3 | 4 | 2 | 1 | 23 | 46 | 47 | 62 | 188 |
| L400 | 4 | 5 | 1 | 1 | 6 | 24 | 56 | 61 | 158 |
| **Total** | 36 | 22 | 16 | 15 | 95 | 282 | 213 | 287 | 966 |

ECoG data acquisition was performed using a TDT RZ2 real-time processor (Tucker-Davis Technologies, Alachua, FL) in the operating room. Collected ECoG data were amplified, filtered (0.7–800 Hz bandpass, 12 dB/octave rolloff), digitized at a sampling rate of 2034.5 Hz and stored for subsequent offline analysis.

### Analysis

Behavioral performance in the target detection task was described in terms of accuracy (hit rate, expressed as % of detected target stimuli), sensitivity index (d’, calculated as Z_hit rate_ − Z_false alarm rate_, where Z(*p*) is the inverse of the cumulative distribution function of the normal distribution) and reaction times (RTs). Only button presses that occurred between the onset of the 5th vowel and the onset of the next trial (i.e. ∼900 ms after the 5th vowel onset; see Fig. 1) were counted as hits. Button presses that overlapped with the next non-target trial were counted as false alarms. The behavioral results thus likely somewhat underestimated the actual target detection rate and biased the RTs towards faster responses.

ECoG data obtained from each recording site were downsampled to 1000 Hz. To minimize contamination from power line noise, ECoG waveforms were de-noised using a demodulated band transform-based procedure (Kovach and Gander, 2016). Data analysis was performed using custom software written in MATLAB Version 7.14 programming environment (MathWorks, Natick, MA, USA).

Two signals of interest were extracted from the recorded data: averaged evoked potentials (AEPs) and high gamma event-related band power (ERBP). AEPs are dominated by low frequency signals, most notably synaptic currents flowing in the vicinity of the electrode, and thus roughly index inputs to the cortical site from which recordings are made. High gamma ERBP is closely related to unit activity and thus largely reflects the output signal from the recorded region (Mukamel et al., 2005; Nir et al., 2007; Ray et al., 2008; Steinschneider et al., 2008; Whittingstall and Logothetis, 2009). Voltage deflections greater than 5 standard deviations from the across-block mean for each recording site were considered artifacts; trials containing such deflections were excluded from further analysis. In 95% of recording sites, 62 or fewer trials (i.e. ≤15.5%) were rejected using this approach.

High gamma power was calculated for each recording site by bandpass filtering the ECoG signal (300th order finite impulse response filter, 70-150 Hz passband), followed by Hilbert envelope extraction. Power envelope waveforms were then log-transformed and, for each of the four sequences within the experimental block (see Fig. 1b), normalized to the mean power over the entire duration of the sequence. LFP and high gamma ERBP waveforms were smoothed using a 4th order Butterworth lowpass filter (30 Hz cutoff). AEPs and high gamma ERBP representing responses to standard and deviant stimuli were computed by time-domain averaging of single-trial LFP waveforms and high gamma ERBP envelopes.

Recording sites that responded to the first four vowels were identified based on the approach used previously by Nourski et al. (2014). First, data within each trial were baseline-corrected by subtracting the mean value in the 100 ms interval immediately preceding the onset of the first vowel. For AEP data, sites were considered responsive to the stimulus onset if either the upper or the lower bound of the 95% confidence interval of the AEP mean was below or above zero µV, respectively, for at least 30 ms, and the following peak exceeded the mean value at the threshold-crossing time point by at least twofold. For high gamma data, the threshold criterion was based on the lower bound of the 95% confidence interval exceeding 0 dB for at least 30 ms, and the following peak exceeded the mean value at the threshold-crossing time point by at least twofold.

LD and GD effects were defined as significant increases in averaged responses to the corresponding deviant vs. standard stimuli within the time interval between 0 and 800 ms following the onset of the fifth vowel. Statistical significance was established using a non-parametric cluster-based permutation test introduced by Maris and Oostenveld (2007). The test statistic was based on grouping adjacent time points that exhibited a significant difference between experimental conditions. The cluster statistic was constructed by first computing two-sample *t*-statistics across all time points for each recording site. For each time point, *t*-values were compared to a threshold corresponding to the 1st percentile tail of the *T*-distribution. The threshold was the 99.5th percentile for two-tailed tests for AEP data and 99th percentile for the one-tailed tests for high gamma data (one-tailed, as high gamma LGD effects were defined as increases in high gamma ERBP in the deviant vs. standard condition). Clusters were defined as consecutive time points for which the *t-*statistic exceeded the threshold. The cluster-level statistic was computed as the sum of the *t*-values within each cluster. The significance level (*p*-value) of those statistics was calculated using permutation tests. To construct the permutation distribution, 10,000 random partitions of experimental conditions were made, shuffled with respect to experimental conditions (standard or deviant), the cluster statistics were calculated, and the largest cluster-level statistic was identified for each partition. This yielded a 10,000-sample distribution set of the test statistics. Monte Carlo *p*-values (Phipson and Smyth, 2010) were calculated for each cluster based on this permutation distribution. The rationale for the maximum cluster-level statistic was to reduce the false alarm rate and control for multiple comparisons at a single-contact level. To correct for multiple comparisons across recording sites, all *p*-values were adjusted by controlling the false discovery rate (FDR) (Heller et al., 2006). FDR correction was applied to all the *p*-values in each of the four test categories (LD AEP, LD high gamma, GD AEP, GD high gamma) using the method introduced by Benjamini and Hochberg (1995). The differences between responses to standard and deviant stimuli were considered significant at *q*<0.05.

Recording sites with at least one significant cluster were considered as exhibiting the deviance effect. The time course of deviance effects was described in three ways. First, onset latencies of deviance effects were defined as the first time point of the earliest significant cluster in each recording site. Second, the detailed time course of each deviance effect was characterized by summing significant clusters across recording sites in all subjects at each time point between 0 and 800 ms following the fifth vowel onset. Third, sites that exhibited LGD effects at four representative time points (100, 225, 400 and 700 ms after the onset of the fifth vowel onset) were plotted in MNI coordinate space.

Differences in onset latencies of LGD effects were evaluated using non-parametric statistical analysis (Kruskal-Wallis tests and two-tailed Wilcoxon rank sum tests). For sites that exhibited both LD and GD effects, comparisons between the effects’ onset latencies were made using two-tailed Wilcoxon signed rank tests.

## Results

### GD stimulus target detection task performance in awake subjects

Subjects exhibited variable performance on the GD stimulus target detection task (Fig. 2a). Target hit rates, computed over the course of the entire block, varied from 51.4% (L372) to 90.3% (R369) (Fig. 2b, upper panel), while sensitivity (*d’*) varied from 1.62 in subject L400 to over 5 in subject R369 (who had no false alarm responses) (Fig. 2b, middle panel). The grand median RT across subjects was 463 ms (Fig. 2b, lower panel).

**Figure 2.**
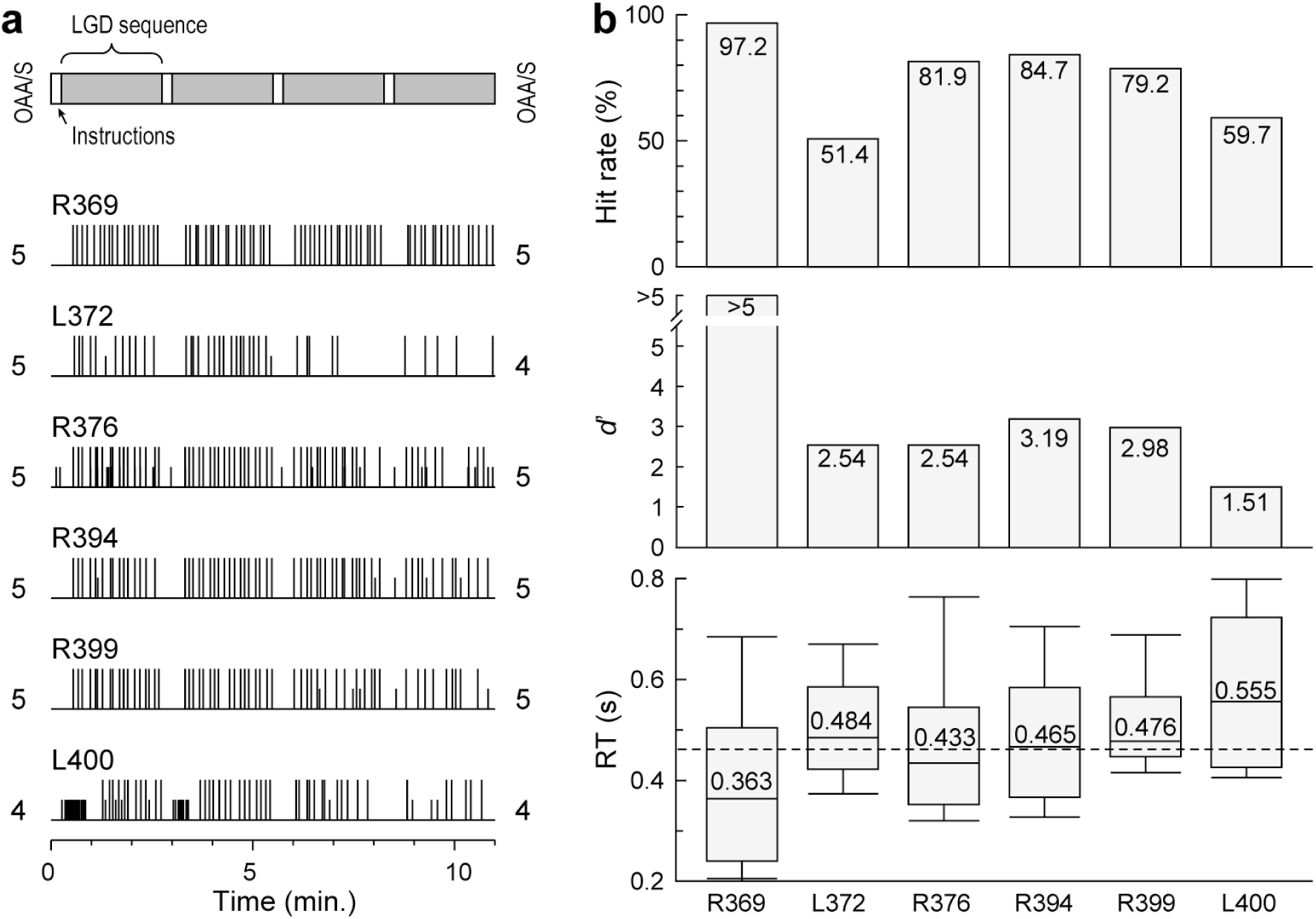
GD stimulus target detection task performance. Summary of data from six subjects. **a:** Timing of button presses in each subject. Hits and false alarm responses are shown as long and short vertical lines, respectively. Schematic of the 11 minute-long experimental block is shown on top; recorded instructions and LGD stimulus sequences (see Fig. 1b) are represented by white and gray rectangles, respectively. Numbers represent OAA/S scores, as assessed before and after the experimental block. **b:** Hit rates (% correctly detected target stimuli), sensitivity (*d*’) and reaction times (RTs) (upper, middle and lower panel, respectively) for each of the six subjects. Box-and-whiskers RT plot depicts median values and 10th, 25th, 75th and 90th percentiles; median values for each subject are shown inside boxes. Dashed line corresponds to the grand median value across all subjects and hit trials (0.463 s).

Four subjects out of six (R369, R376, R394, R399) had maximal OAA/S scores of 5 before and after completion of the experimental block, indicative of a fully awake and alert state. Task performance for subject L372 was characterized by a relatively low hit rate, yet the second highest sensitivity. Analysis of this subject’s task performance over the course of the experimental block showed a marked decrease in button presses indicative of decreased attention to the task (see Fig. 2a). This paralleled the decrease in that subject’s level of awareness as indexed by OAA/S score. Thus, within the block, when the subject was alert and responding, performance accuracy was high. Likewise, the baseline level of alertness in subject L400 was diminished compared to the rest of subjects, as evidenced by OAA/S scores. Thus, as expected, task performance was related to the overall behavioral state of the subjects.

### Neural responses to the vowels prior to the onset of deviance

The first four vowels of the experimental five-vowel stimuli occurred prior to the emergence of either local or global deviance. Examination of neural activity during this initial portion of the stimuli provided a means to examine responses elicited by sound, regardless of auditory novelty. The vowel tokens elicited AEPs and high gamma responses in all ROIs (Fig. 3). Responses simultaneously recorded from four exemplary right hemisphere sites (Fig. 3a) in subject R369 who had both the highest hit rate and sensitivity in the behavioral task are shown in Figure 3b. Responses to the first four vowels varied as a function of ROI. Within HGPM (Fig. 3b, upper row), AEP waveforms were time-locked to the onset of each vowel. High gamma increases were primarily associated with the onset of the five-vowel stimulus. AEPs and high gamma responses were of lower magnitude in other ROIs (Fig. 3b, second – fourth rows). The overall incidence of significant AEPs (see Methods) varied across cortical regions. Specifically, across all six subjects, 100% sites in HGPM, HGAL and PT, 60.0% in PP, 81.1% in STG, 48.2% in auditory-related cortex and 35.7% in PFC exhibited significant AEPs to the first four vowels. High gamma activation was more spatially restricted; significant responses were present in 94.0% of HGPM sites, 54.5% in HGAL, 62.5% in PT, 13.3% in PP, 40.0% in STG, 4.96% in auditory-related cortex and 0% in PFC.

**Figure 3.**
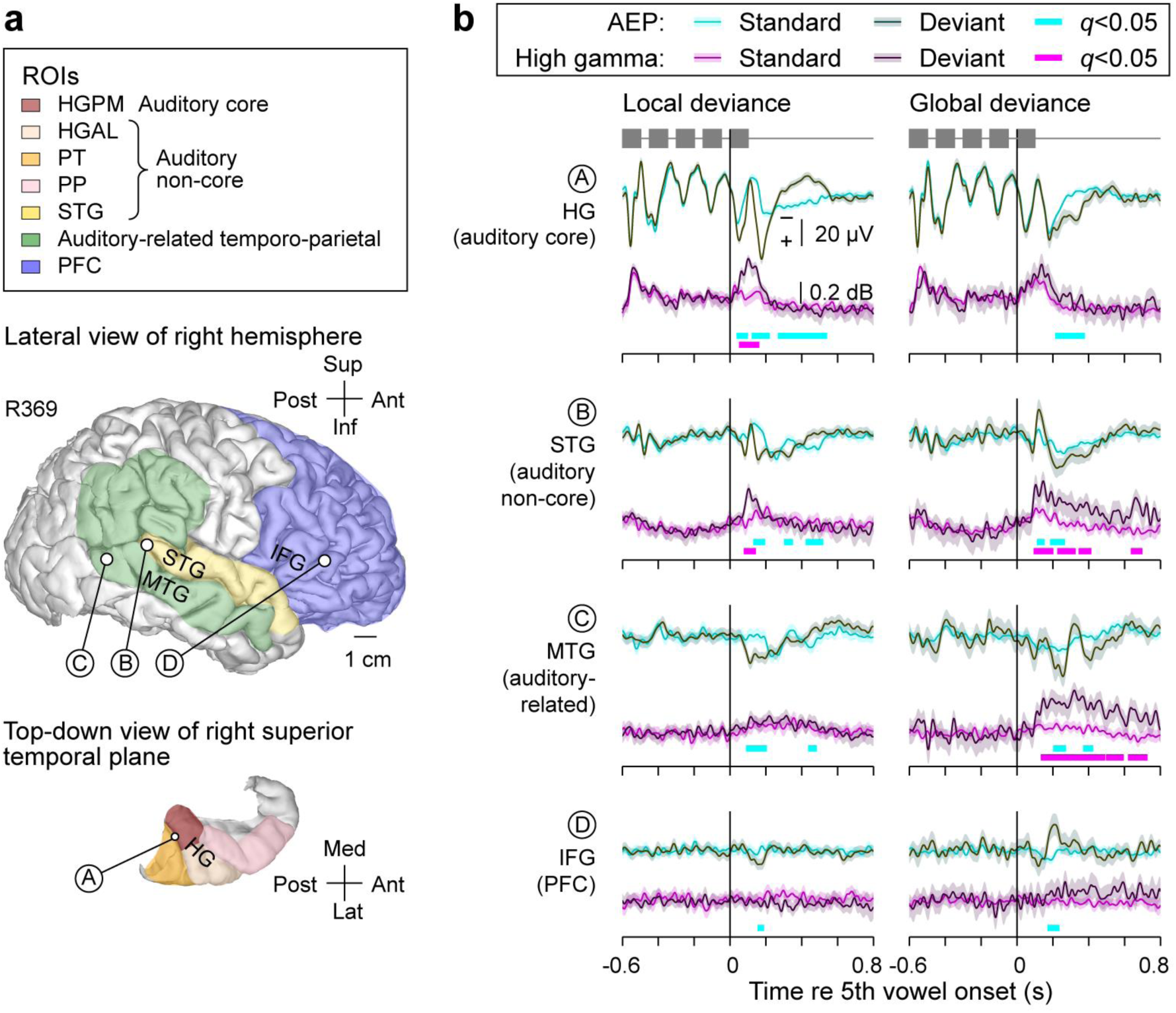
Cortical responses to standard and deviant stimuli along the auditory processing hierarchy. **a:** MRI top-down view of superior temporal plane and side view of the hemispheric surface showing the locations of four representative recording sites (sites A, B, C, D in HG, STG, MTG and IFG, respectively) in subject R369. Colors represent different ROIs used in the present study. **b:** AEP and high gamma responses recorded from the four cortical sites in response to standard and deviant stimuli (blue and red waveforms, respectively). Lines and shading represent mean values and the 95% confidence intervals, respectively. LD and GD effects are presented on the left and right, respectively. Green bars highlight significant differences between responses to standard and deviant stimuli (*q*<0.05, non-parametric cluster-based permutation test, FDR-corrected).

### Spatial properties of LD and GD effects

Auditory novelty elicited AEPs and increases in high gamma ERBP beyond those that might represent offset responses to the final token. These deviance effects were defined as significant differences between responses to standard and deviant stimuli (cyan and magenta bars in Fig. 3b). In subject R369, significant LD effects were seen in the AEP at all ROIs, while high gamma increases were restricted to core and non-core auditory cortex (see Fig. 3b, left column). GD AEP effects were also present at all four sites, with the earliest changes occurring in the STG, followed by activity in PFC. Significant high gamma GD effects were present in the STG and auditory-related cortex in the MTG (see Fig. 3b, right column).

Exemplar data are representative of the overall spatial distribution of LD and GD effects in this subject (Fig. 4). AEPs reflecting LD were seen throughout all ROIs as well as other brain regions (e.g. amygdala; inset in Fig.4a), whereas increases in high gamma ERBP were restricted to auditory cortex ROIs (Fig. 4a). GD effects were associated with a different spatial distribution compared to LD effects (Fig. 4b). High gamma GD effects were not common in auditory cortex and instead were more prominent in auditory-related cortex and PFC, reflecting activity originating within higher levels of the cortical processing hierarchy. Comparable patterns were observed in subject R376 (Supplementary Fig. 2). Especially notable was the prevalence of LD, but not GD high gamma effects within superior temporal plane, and the opposite pattern (GD, but not LD high gamma effects) in PFC.

**Figure 4.**
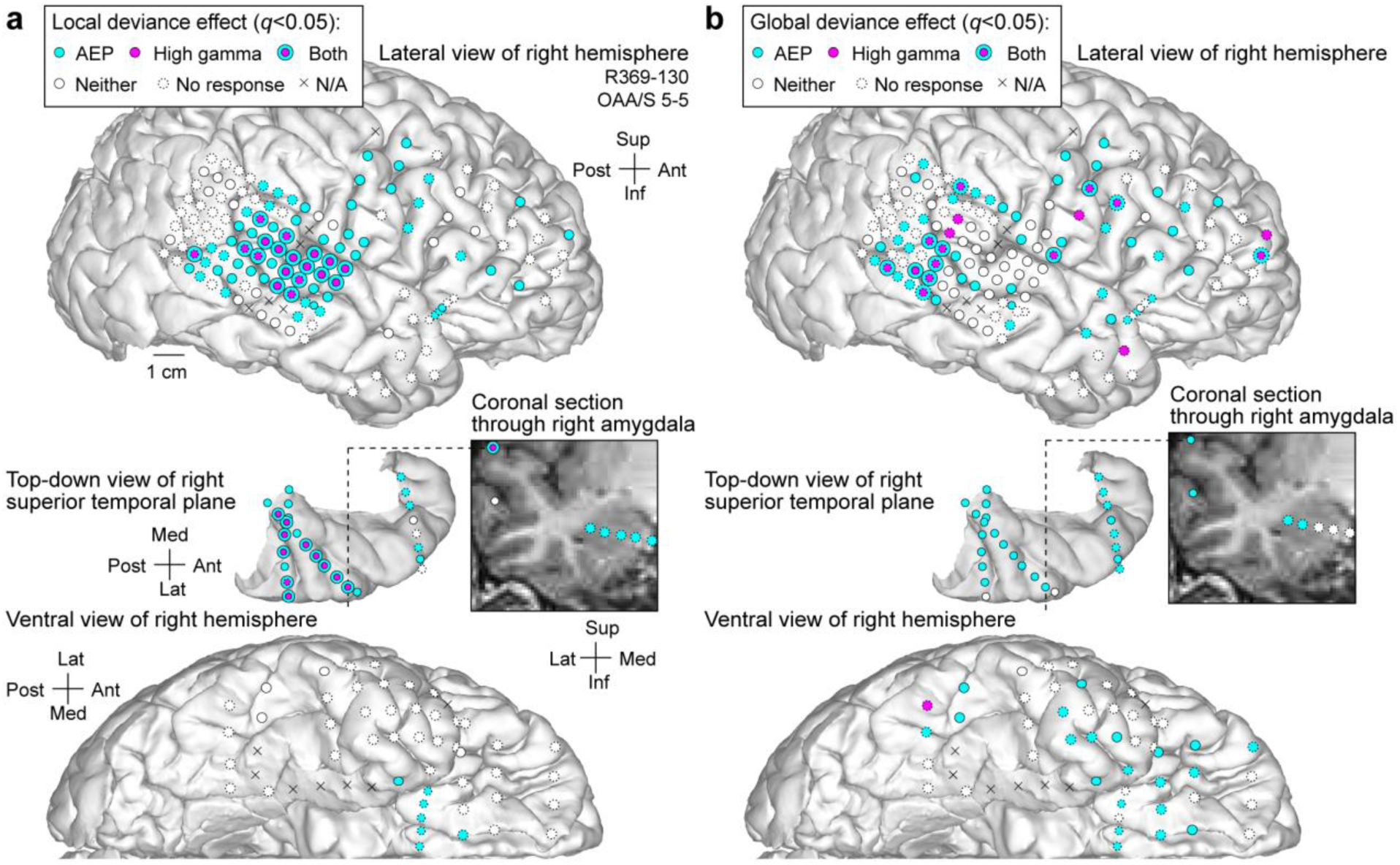
Spatial distribution of sites exhibiting LD and GD effects (panels **a** and **b**, respectively) in a representative subject (R369). Sites that feature significant differences between responses to standard and deviant stimuli, as measured by AEP and high gamma responses, are shown in cyan and magenta, respectively. Sites that didn’t exhibit either significant difference are shown in white. Dashed outlines indicate absence of response to the initial portion of the stimuli. Sites that were not included in the analysis are marked with X. **Insets:** MRI coronal section through the right temporal lobe (section plane indicated by a dashed line) showing location of depth electrode contacts in the right amygdala that exhibited AEP local and deviance effects.

Across the entire subject cohort, there was a substantial spatial overlap in LD and GD effects, consistent with involvement of many sites in both LD and GD networks (Fig. 5). AEP LD effects were prominent in the superior temporal plane and were usually associated with GD effects (Fig. 5a). In contrast, sites exclusively responsive to GD were rare here. Beyond the superior temporal plane, AEP LD effects were widely distributed throughout all ROIs and often co-located with GD effects. Sites exclusively exhibiting AEP GD effects were rare in auditory cortex on the STG, yet prevalent in surrounding areas, including MTG, SMG, angular gyrus, as well as PFC.

**Figure 5.**
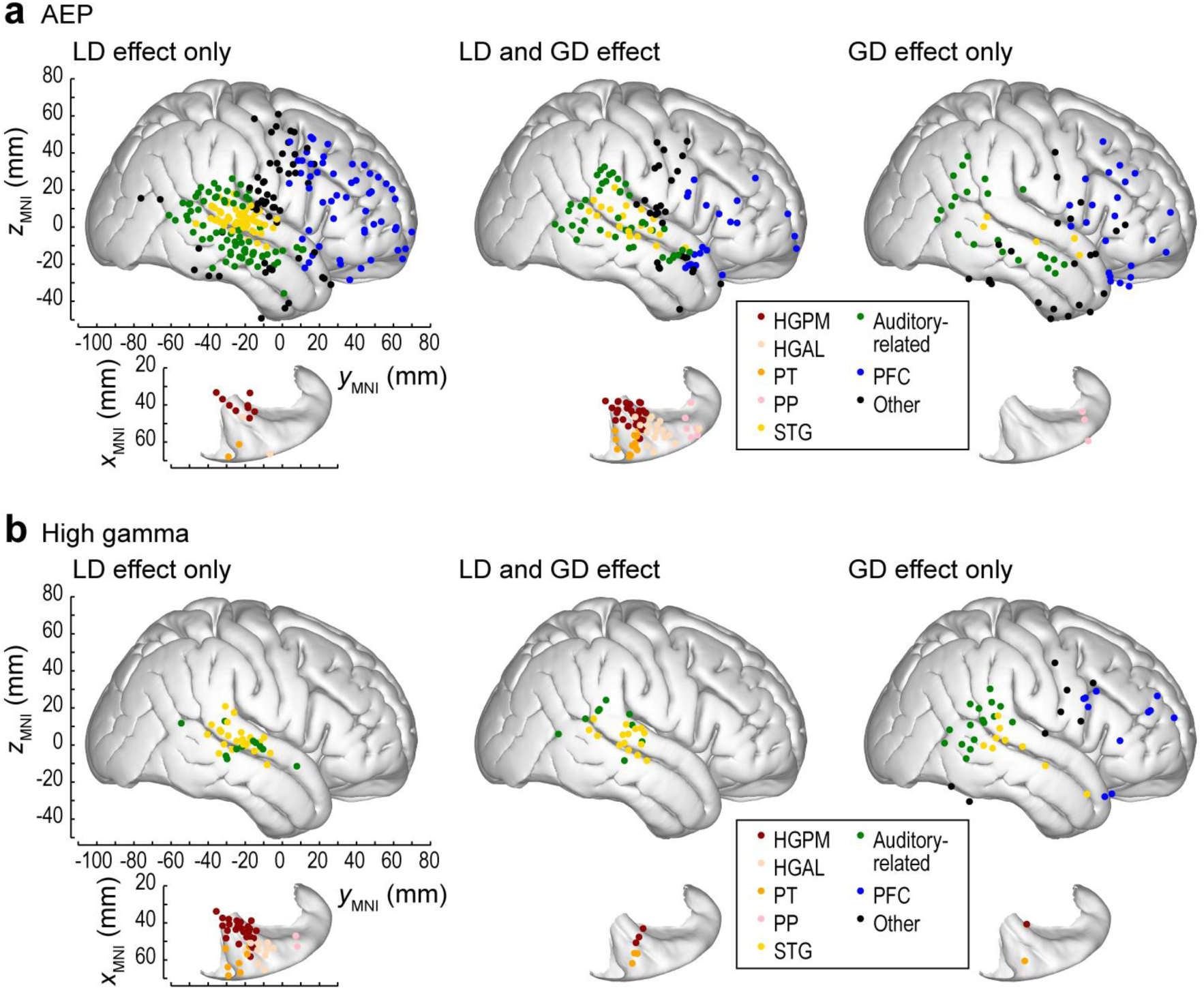
Spatial distribution of sites exhibiting significant LD and GD effects as measured by the AEP and high gamma activity (panels **a** and **b**, respectively). Summary of data from six subjects, plotted in MNI coordinate space and projected onto FreeSurfer average template brain. Top-down views of the right superior temporal plane are plotted underneath side views of the right lateral hemispheric convexity, ^aligned with respect to the *y*^MNI ^coordinate. Sites exhibiting significant LD effects only, both LD and GD^ effects, and GD effects only, are depicted in the left, middle and right column, respectively. Sites are color-coded based on their ROI assignment in each individual subject.

A sparser representation of LGD effects was seen in high gamma activity (Fig. 5b). In contrast to AEP, sites in the superior temporal plane largely exhibited only high gamma LD effects, and not GD effects. High gamma GD effects (with or without LD effects) were more prevalent in the STG (25 out of 95 STG sites) compared to all other ROIs. Within the PFC, high gamma GD effects, but not LD effects, were observed.

### Temporal properties of LD and GD effects

Onset latencies (i.e. the timing of the first significant difference after the fifth vowel stimulus) of the LD effects were shortest in auditory cortex located in the superior temporal plane (HGPM, HGAL and PT), and became progressively longer in non-core auditory cortex on the lateral STG, auditory-related cortex and PFC (Fig. 6a, left column). AEP and high gamma onset latencies within the superior temporal plane were comparable between core (HGPM) and non-core regions (HGAL, PT) (AEP: *p* = 0.986, high gamma: *p* = 0.714; Kruskal-Wallis test). In contrast, onset latencies of AEP LD effects increased from HGPM to non-core auditory cortex on the lateral STG, from STG to auditory-related cortex, and from auditory-related cortex to PFC (HGPM vs. STG *p* = 0.0229, *z* = −2.28, *W* = 1585; STG vs. auditory-related *p* = 0.001006, *z* = −3.29, *W* = 5321; auditory-related vs. PFC *p* = 0.00155, *z* = −3.17, *W* = 9655; two-tailed Wilcoxon rank sum tests). Likewise, onset latencies of high gamma LD effects increased from HGPM to STG (*p* = 0.00382, *z* = −2.89, *W* = 724), although no significant difference was found between STG and auditory-related cortex (*p* = 0.176, *z* = −1.35, *W* = 1085; two-tailed Wilcoxon rank sum tests), and high gamma LD effects were absent in PFC. The relatively low occurrence of LGD effects within PP (LD AEP: 6 sites out of 15 across 5 subjects; LD high gamma: 2 sites; GD AEP: 9 sites; GD high gamma: 0 sites) precluded comparisons of LGD effect onset latencies in this region with other ROIs.

**Figure 6.**
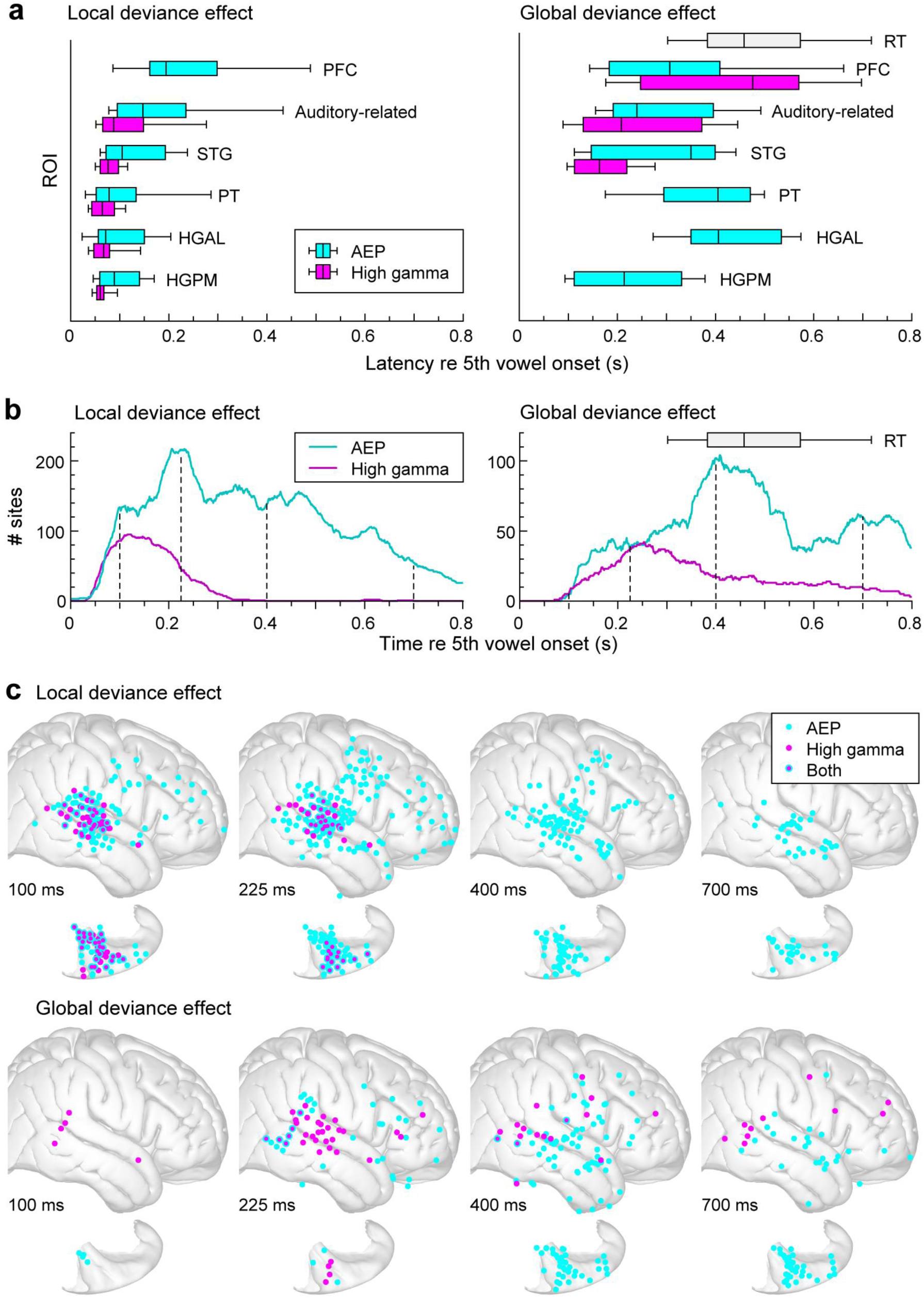
Temporal properties of LD and GD effects. **a:** Onset latencies of LD and GD effects (left and right column, respectively) across ROIs. Summary of data from six subjects. Onset latency distributions of AEP and high gamma effects are shown in cyan and magenta, respectively. Box and whiskers plots are shown for ROIs with at least ten sites across the six subjects exhibiting significant deviance effects and show across-contact medians, quartiles, 10th and 90th percentiles. Gray box represents RTs across the six subjects (see Fig. 2b for measurements in individual subjects). **b:** Time course of LD and GD effects (left and right column, respectively). Total number of sites across all subjects exhibiting significant differences between responses to standard and deviant stimuli is plotted as a function of time after the 5th vowel onset for AEP and high gamma in cyan and magenta, respectively. Vertical dashed lines represent four exemplar time points depicted in panel c. **c:** Spatial distribution of LD and GD effects (upper and lower row) at four representative time points (left to right: 100, 225, 400 and 700 ms after 5th vowel onset). Sites that exhibited significant deviance effects at these time points are shown in cyan and magenta for AEP and high gamma, respectively. Summary of data from six subjects, plotted in MNI coordinate space and projected onto FreeSurfer average template brain.

A markedly different latency profile was seen for GD effects (Fig. 6a, right column). First, latencies were generally longer compared to LD effects (Table 3). A total of 176 sites exhibited both LD and GD AEP effects, while 35 sites were characterized by presence of both LD and GD high gamma effects. For these sites, pairwise comparisons revealed that LD effects were significantly earlier (AEP: *p* = 2.84×10^−14^, *z* = −7.61, *W* = 2640; high gamma: *p* = 6.645×10^−6^, *z* = −4.51, *W* = 40; two-tailed Wilcoxon signed rank tests).

**Table 3.** Comparison of LD and GD effect onset latencies across ROIs.

| ROI | AEP LD vs. GD |  |  |  |  | High gamma LD vs. GD |  |  |  |  |
| --- | --- | --- | --- | --- | --- | --- | --- | --- | --- | --- |
| | LD effect | | GD effect | | LD Vs. GD $p$-value | LD effect | | GD effect | | LD vs. GD $p$-value |
| | Median lat. (ms) | $n_{sites}$ | Median lat. (ms) | $n_{sites}$ | | Median lat. (ms) | $n_{sites}$ | Median lat. (ms) | $n_{sites}$ | |
| HGPM | 88.5 | 36 | 214 | 27 | $2.89 \times 10^{-5}$ | 57 | 28 | 192 | 4 | **0.00333** |
| HGAL | 71 | 21 | 406 | 19 | $1.71 \times 10^{-7}$ | 64 | 11 | - | - | - |
| PT | 78 | 16 | 405 | 14 | $2.04 \times 10^{-5}$ | 61 | 10 | 215.5 | 4 | **0.00199** |
| PP | 217.5 | 6 | 353 | 9 | 0.627 | 143 | 2 | - | - | - |
| STG | 104.5 | 70 | 350 | 25 | $3.31 \times 10^{-7}$ | 73 | 39 | 163.5 | 24 | $4.44 \times 10^{-9}$ |
| Auditory-related | 147 | 114 | 240 | 64 | $2.75 \times 10^{-8}$ | 84.5 | 20 | 208 | 25 | $8.14 \times 10^{-5}$ |
| PFC | 194 | 75 | 307 | 44 | **0.00326** | - | - | 476 | 11 | - |

High gamma GD effects were relatively uncommon in the superior temporal plane (4/36 sites in HGPM, 0/22 in HGAL, 4/16 sites in PT and 0/15 in PP), precluding statistical analyses of onset latencies in these ROIs. Of the remaining ROIs, median latencies of high gamma GD effects were shortest in lateral STG, though they did not differ significantly from those in auditory-related cortex (STG vs. auditory-related *p* = 0.0854, *z* = −1.72, *W* = 513.5). Median latencies of high gamma GD effects were longest in PFC (STG vs. PFC *p* = 7.43×10^−5^, *z* = −3.96, *W* = 320; auditory-related vs. PFC: *p* = 0.00251, *z* = −3.02, *W* = 374).

The paucity of high gamma GD effects in HGPM, HGAL, PT and PP contrasted with presence of AEP GD effects. Onset latencies of AEP GD effects were different across ROIs (*p* = 3.91×10^−5^; Kruskal-Wallis test). Interestingly, AEP GD onset latencies were the shortest in HGPM (*p* < 0.05 in all pairwise comparisons; two-tailed Wilcoxon rank sum tests), even though the shortest-latency high gamma GD effects were found on STG, and high gamma GD effects were sparse in core auditory cortex. High gamma GD effects preceded AEP GD effects in all ROIs except HGPM (HGPM: *p* = 0.304, *z* = −1.03, *W* = 569; *p* < 0.05 for pairwise comparisons in all other ROIs; two-tailed Wilcoxon rank sum tests). Finally, high gamma activity in STG and auditory-related cortex preceded the subjects’ behavioral responses (STG: *p* = 3.65×10^−14^, *z* = −7.57, *W* = 590; auditory-related: *p* = 6.89×10^−10^, *z* = −6.17, *W* = 1387), while high gamma GD effect onset latencies in PFC overlapped with the subjects’ reaction times (*p* = 0.809, *z* = −0.242, *W* = 1787).

Overall timing of LD and GD effects provides evidence for region- and effect-specific activation of underlying generators. AEP LGD effects had a more extended and complex time course compared to high gamma effects (Fig. 6b). Figure 6b shows the estimated overall probability density function of cortical activation during LD and GD effects, computed by summing across subjects the number of sites exhibiting significant effects at each time point relative to the onset of the fifth vowel. AEP LG and GD effects were characterized by multi-peaked time courses, while high gamma effects exhibited simpler time courses. AEP LD effects were most prominent at 200-250 ms after the fifth vowel onset, while AEP GD effects were dominated by a peak at 400 ms. The overall time course of the high gamma LD effect was characterized by a single peak around 125 ms after the fifth vowel onset, while the high gamma GD effect peaked at 250 ms.

The spatiotemporal evolution of LGD effects is summarized in Figure 6c for four exemplar time points (see dashed lines in Fig. 6b). Early LD effects were represented by distributed AEPs in all ROIs and high gamma effects in all ROIs except PFC. Over time, LD effects became progressively restricted to AEPs within temporal cortex. GD effects were more sparsely represented, and featured high gamma activity that developed at longer latency and was more broadly distributed across all time points. Taken together, these results demonstrate that LD and GD effects are distinct both in their spatial and temporal profiles.

## Discussion

### Summary of findings

The data reveal distinct spatial and temporal profiles of neural responses underlying auditory novelty detection at short (LD effect) vs. long (GD effect) time scales. Both effects, as indexed by AEPs, were broadly distributed across auditory, auditory-related and prefrontal cortical regions. In contrast, high gamma LD effects were focused mainly in auditory core and non-core areas, while high gamma GD effects were more concentrated in auditory-related cortex and PFC. The regional distribution of latencies of LD effects can be interpreted within the context of the predictive coding model. Latencies are earliest within core and non-core regions of auditory cortex and become progressively longer along the processing hierarchy. This pattern is consistent with FF signal propagation of prediction errors. By contrast, the shortest latencies of high gamma GD effects were observed in STG and auditory-related cortex, and preceded the onset of AEP GD effects both within those areas and in other ROIs. Thus, signal propagation, and presumably the composition of the network driving the predictive coding process, differs for LD versus GD effects. The latter appear to be initiated in higher order regions of auditory cortex and in auditory-related temporoparietal cortex rather than in PFC. These findings help clarify the involvement of specific brain regions in the predictive coding network subserving auditory novelty detection.

### Cortical generators of LD and GD effects

While auditory cortex has been consistently observed to contribute to LD effects, there is conflicting evidence for the additional involvement of frontal cortical generators. Some studies reported that LD effects are confined to temporal cortex (Baudena et al., 1995; Bekinschtein et al., 2009; El Karoui et al., 2015), whereas others have also shown involvement of PFC using ECoG (Liasis et al., 2001; Rosburg et al., 2005) as well as EEG/MEG and fMRI (reviewed in Deouell, 2007). The data presented here clearly indicate widespread cortical activation associated with LD effects: AEPs were observed in all ROIs, including parietal and frontal cortex, and high gamma signals observed in all areas except PFC. These data suggest involvement of multiple levels of the auditory cortical hierarchy, though the absence of high gamma LD effects in PFC suggests a passive role of the frontal generator for novelty detection over this timescale.

We observed prominent GD effects (AEPs and high gamma) in STG, auditory-related cortex and in PFC, consistent with previous reports (Bekinschtein et al., 2009; Chennu et al., 2013; El Karoui et al., 2015). Although there was a substantial spatial overlap of AEP GD and LD effects, recording sites exhibiting only GD effects were largely outside of auditory cortex ROIs. The spatial distributions of high gamma GD and LD effects exhibited less overlap than for AEPs, suggesting that the populations of cells initiating these effects are largely distinct, with a common node in both networks located in STG.

### Auditory novelty detection over multiple time scales

There are multiple electrophysiological signatures of auditory novelty detection, distinct in their generators, latency, and dependence on brain state (Friedman et al., 2001; Kok, 2001; Naatanen and Alho, 1995; Naatanen et al., 2011). At a mechanistic level, responses to auditory novelty within auditory cortex can be ascribed in part to stimulus-specific adaptation (Eliades et al., 2014; Fishman and Steinschneider, 2012; Ulanovsky et al., 2003). This process, in which the long-term statistics of preceding sounds modulate responses to subsequent sounds in a context-dependent manner, likely contributes to both LD and GD effects within auditory cortex (Holt, 2006; Ulanovsky et al., 2004). However, modeling studies (Wacongne et al., 2012), as well as evidence from omission responses (Wacongne et al., 2011) and from MMNs generated in response to abstract rule violations (Paavilainen, 2013), suggest that higher order and especially FB-mediated processes are also involved in these novelty signals.

While it is necessary to be cautious when comparing intracranial data with scalp-recorded potentials or MEG fields, it is noteworthy that the AEP LD effect reported here was prominent at 150-250 ms after the fifth vowel onset. This timing is consistent with previous reports identifying the LD effect with the MMN (Bekinschtein et al., 2009; Wacongne et al., 2011). More broadly, onset latencies of AEP LD effects across ROIs had a wide range (50 – 500 ms; see Fig. 6a), comparable to that reported in a previous intracranial study (70 – 440 ms; El Karoui et al., 2015), and overall time course of AEP LD effects extended over nearly the entire 800 ms analysis window (Fig. 6b). Such a wide range of latencies is consistent with involvement of a broad network in detection of local deviance. Indeed, AEP LD effects were observed in all ROIs investigated. Thus, the present findings are in disagreement with previous studies that reported focal expression of the LD effect in STG (Bekinschtein et al., 2009; Liebenthal et al., 2003; Sabri et al., 2004; Wacongne et al., 2011), but are consistent with studies that reported LD effects within PFC (Durschmid et al., 2016; Rinne et al., 2000; Uhrig et al., 2016). Some of these disparities may result from differences in signal-to-noise ratios in the recordings. LD effects were prominent and of large amplitude in auditory cortex and adjacent regions, and it is these areas where differences in responses between standard and deviant stimuli are most likely to reach statistical significance.

In the current study, a progressive increase in latency of both AEP and high gamma responses was observed along the ascending hierarchy of the ROIs exhibiting LD effects (see Fig. 6a). (Durschmid et al., 2016). This latency profile likely includes FF signal propagation during deviance detection. However, the temporally and spatially distributed response profiles suggest mechanisms more complex than a simple FF process. The range of latencies in each ROI, especially the longer latencies of AEPs relative to high gamma ERBP, and the extended overall time course of LD effects, are consistent with bidirectional signal flow in a broad network operating over multiple spatial and temporal scales, part of which is automatic/pre-attentive, and part of which is active/predictive (El Karoui et al., 2015; Naatanen et al., 2011; Sculthorpe et al., 2009; Strauss et al., 2015)

Unlike the LD effect, the GD effect is dominated by response components with onset latencies greater than 200 ms, and has been associated with the P3b component of the event-related potential (Bekinschtein et al., 2009; Wacongne et al., 2011), a task- and attention-dependent index of contextual updating that operates over time scales of seconds to minutes (Kok, 2001; Polich, 2007). In the present study, the latency distributions for LD and GD effects overlapped (see Fig. 6a), but the mean latencies of GD effects were significantly longer than those of LD across a wide range of recording sites (see Table 3). Previous reports have emphasized a distributed network engaged by global deviance detection, involving anterior cingulate, parietal, temporal and prefrontal regions (Bekinschtein et al., 2009; El Karoui et al., 2015; Uhrig et al., 2014). This overlaps with the postulated global workspace network subserving conscious sensory processing (Dehaene and Changeux, 2011; Dykstra et al., 2017). Similar to these studies, the present study revealed a GD network widely distributed in space and time, with significant responses observed in all regions of interest, from core auditory cortex to PFC (Fig. 5), and over the time window 100 – 800 ms after the onset of the fifth vowel (Fig. 6b). The spatial overlap of LD and GD networks may either be representative of a commonality in mechanisms underlying the two effects (e.g. forward masking at multiple time scales) or the co-location of two distinct processes that remain to be elucidated (see Fig. 5).

The shortest onset latencies of GD effects corresponded to high gamma ERBP in the STG and adjacent auditory-related cortex. By contrast, high gamma GD effects were uncommon in the superior temporal plane ROIs (HGPM, HGAL, PT and PP), compared to reliably observed AEP GD effects (Fig. 5). Given that high gamma is thought to represent a population-level surrogate for unit activity (Mukamel et al., 2005; Nir et al., 2007; Steinschneider et al., 2008), the onset latency data support a model in which GD effects trigger earliest spiking activity in STG and auditory-related cortex, and signals propagate both down the hierarchy via FB projections triggering AEPs in lower areas and up the hierarchy to PFC via FF projections where they trigger both AEPs and spiking activity. This is in contrast to models in which long time scale predictive coding relies on FB projections from frontal cortex (cf. Durschmid et al., 2016), and may be relevant to the current debate over the importance of frontal versus parietal regions for the neural basis of consciousness (Boly et al., 2017; Odegaard et al., 2017; Siclari et al., 2017). Functional connectivity analyses may be necessary to further identify specific sources of GD effects.

At present, it is unclear to what degree GD effects occur independently from LD effects in STG. Emergence of GD effects in auditory-related cortex can occur in the absence of LD effects (see Fig. 5). This observation suggests that temporoparietal auditory-related cortex is operating over temporal windows of integration greater than those that characterize auditory cortex. This is parsimonious with regional differences in speech processing at acoustic-phonemic and lexico-semantic levels, wherein the former is processed in STG and the latter in surrounding auditory-related regions (Nourski et al., 2016).

AEP GD effects had relatively short latencies in HGPM (see Fig. 6a). The time course of AEP GD effects in HGPM was similar to the overall time course of AEP GD effects, with the early components most prominent at 100-200 ms followed by components centered at 400 and 700 ms (see Fig. 6b). As noted previously (El Karoui et al., 2015; Wacongne et al., 2011), the early components likely arise due to the partial interdependence of LD and GD effects. This interaction, or context dependence, arises because the magnitude of LD effects is inversely correlated with the probability of the deviant stimulus. Thus, trial sequences in which the LD stimulus is also the GD (see Fig. 1c) will have large, short-latency responses to that stimulus. This may simply reflect the difference in probability of occurrence of the LD stimulus, but could also reflect violation of a second order prediction about the vowel sequence that operates on a longer time scale (Ulanovsky et al., 2004). Modifications of the classic LGD paradigm that include token omission as a means to elicit global deviance may help clarify these outstanding issues (Wacongne et al., 2011).

### Caveats and limitations

Principal caveats of the present study are related to the subject population and limitations of the experimental paradigm. The former is an issue of concern inherent to all human intracranial studies. Experimental subjects have a neurologic disorder, and their brain responses may not be representative of a healthy population (Nourski and Howard, 2015). To address this issue, recordings from epileptic foci in each subject were excluded from the analyses. Most importantly, results were replicated in multiple subjects, who had different neurologic histories, seizure foci and antiepileptic drug regimens. Cognitive function in each subject was in the average range, and all subjects were able to perform the experimental task successfully.

Another potential limitation inherent to studying patients undergoing chronic invasive monitoring is electrode coverage based solely on clinical needs. In general, coverage is unilateral. This issue precludes within-subject comparisons across cerebral hemispheres. Four of the six subjects studied here had coverage of the non-dominant right hemisphere. This raises the question of whether recordings in the non-dominant hemisphere are directly relevant to the experimental task. Results of non-invasive studies including those using fMRI, EEG and MEG all demonstrate bihemispheric processing of auditory novelty (Bekinschtein et al., 2009; Wacongne et al., 2011; Strauss et al., 2015). The relatively poorer task performance of the two subjects with left hemisphere seizure foci and left hemisphere electrode coverage likely reflected their overall lower level of alertness as indexed by OAA/S scores. The extent of electrode coverage and spatial distribution of LGD effects in these two subjects was comparable with the rest of the subject cohort.

Finally, during the target detection task, neural activity associated with the motor act of button press may temporally overlap with responses associated with deviance detection *per se*. This confound was minimized in the present study by having subjects operate the response button with the hand ipsilateral to the hemisphere from which recordings were made (see Nourski et al., 2016, for a further discussion of this issue).

### Functional significance

The LGD experimental paradigm represents an important empirical test of the predictive coding model. First, the first four vowels within an LGD sequence are always the same and thus by themselves do not generate an error signal. Cortical responses to this initial portion of the LGD stimuli become progressively weaker along the ascending auditory hierarchy, in accordance with the predictive coding model. Second, the relative involvement of lower and higher hierarchical stages in processing of local and global deviance, respectively, highlights the utility of the results obtained in this experimental paradigm, and may enhance novelty response generator identification as seen in non-invasive studies (e.g. Symonds et al., 2017). This paradigm has potential use as part of a noninvasive metric of conscious processing. For example, evidence suggests that clinical evaluation alone is insufficient to fully evaluate patients with disorders of consciousness (Bayne et al., 2017; Bernat, 2017; Naccache, 2017; Schnakers et al., 2009). The present study also serves as a foundation for ongoing and future studies that seek to gain better understanding of sensitivity of auditory cortical novelty processing to general anesthesia (Nourski et al., 2018). In turn, knowledge gained from this work will contribute to improvements in accuracy of assessing conscious processing in clinical and non-clinical settings.

## Acknowledgements

We are grateful to Haiming Chen, Phillip Gander, Bradley Hindman, Christopher Kovach, Bryan Krause, Rashmi Mueller, Caitlin Murphy, Yuri Saalmann, Beau Snoad and Xiayi Wang for help with data collection, analysis and comments on the manuscript.

