## Supplementary Materials for "Processing of Auditory Novelty Across the Cortical Hierarchy: An Intracranial Electrophysiology Study"

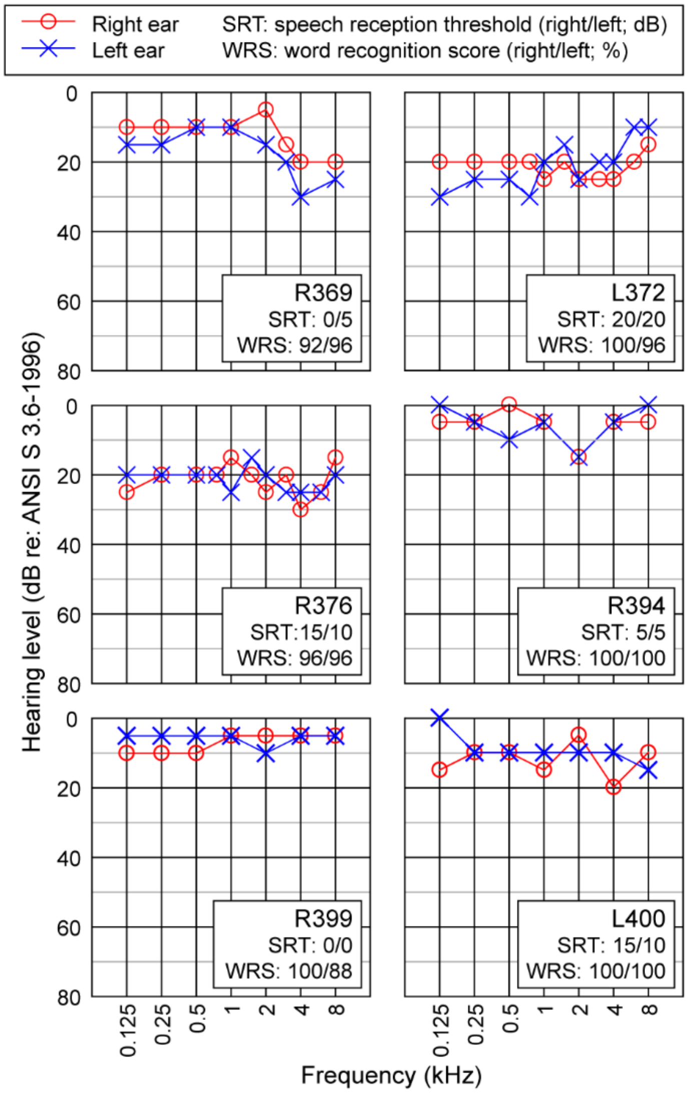


**Supplementary Figure 1.** Audiological evaluation data. Pure-tone thresholds (air conduction) are plotted for right and left ear (red and blue plots, respectively) for each subject. Speech reception threshold and word recognition score values are presented for right and left ear in the Figure legend for each subject.
