## Supplementary Materials for "Processing of Auditory Novelty Across the Cortical Hierarchy: An Intracranial Electrophysiology Study"

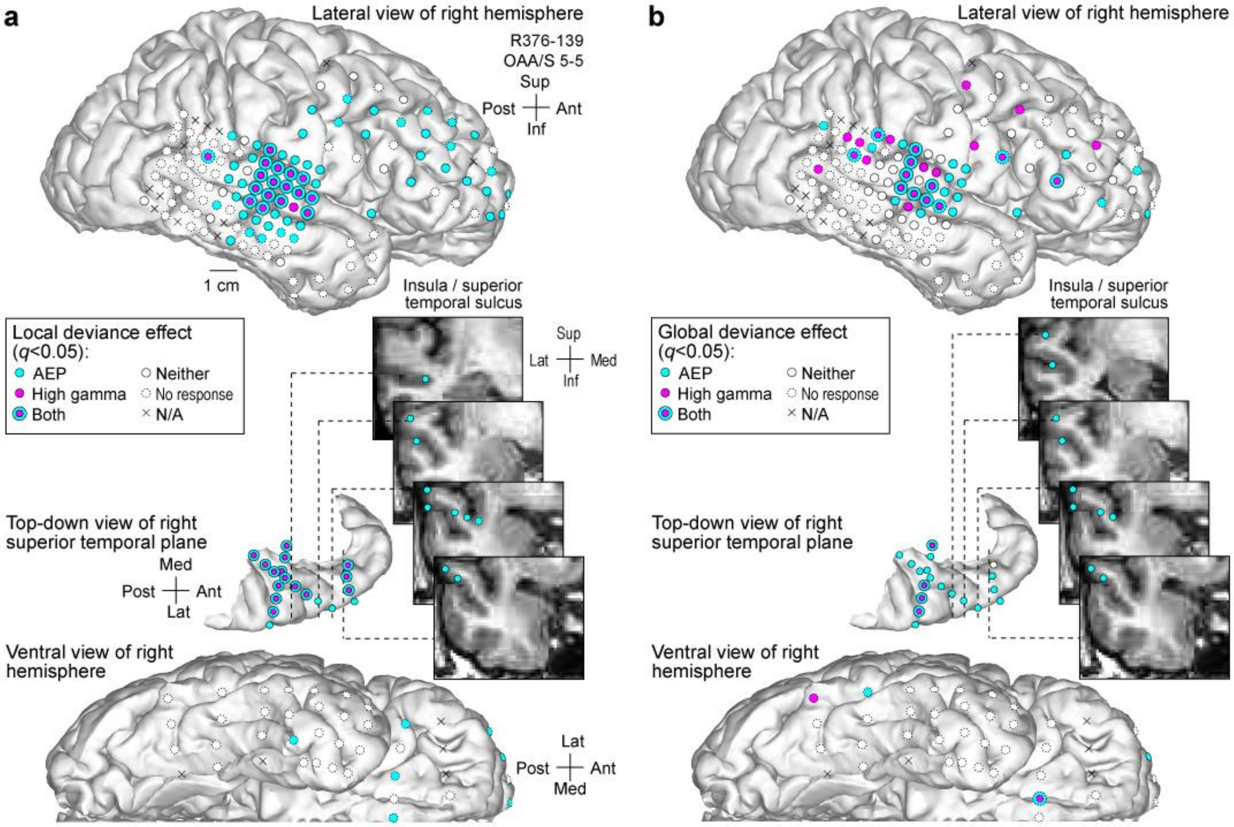


**Supplementary Figure 2.** Spatial distribution of sites exhibiting LD and GD effects. Confirmatory data from subject R376. See legend of Fig. 4 for details. **Insets:** MRI coronal sections through the right temporal lobe (section planes indicated by dashed lines) showing locations of depth electrode contacts in the right insular cortex and superior temporal sulcus that exhibited AEP LD and GD effects.
